# Head-on replication-transcription collisions lead to formation of life threatening R-loops

**DOI:** 10.1101/154427

**Authors:** Kevin S. Lang, Ashley N. Hall, Christopher N. Merrikh, Mark Ragheb, Houra Merrikh

## Abstract

Encounters between transcription and DNA replication machineries lead to conflicts that shape genomes, influence evolution, and lead to genetic diseases in humans. Although unclear why, head-on transcription (lagging strand genes) is especially disruptive to replication, increases DNA breaks, and promotes mutagenesis. Here, we show that head-on replication-transcription conflicts lead to pervasive RNA:DNA hybrid formation in *Bacillus subtilis*. We find that replication beyond head-on conflict regions requires the activity of a RNA:DNA hybrid processing enzyme, RNase HIII. Remarkably, pervasive RNA:DNA hybrid formation at head-on genes completely stops replication and inhibits gene expression in a replication-dependent manner. Accordingly, we find that resolution of head-on conflicts by RNase HIII is crucial for survival upon exposure to various stresses, as many stress response genes are encoded head-on to replication. We conclude that R-loops, RNA:DNA hybrids formed outside of the transcription bubble, exacerbate head-on replication-transcription conflicts, thereby threatening life, especially upon exposure to environmental stresses.

## INTRODUCTION

DNA replication and transcription function at the same time and on the same DNA template. The coupling of these two essential processes leads to conflicts between the two machineries that disrupt DNA replication, lead to genomic rearrangements, cause breaks in the DNA, and increase mutagenesis (French, 1992; Merrikh et al., 2015; Million-Weaver et al., 2015a; Sankar et al., 2016; Srivatsan et al., 2010; Tuduri et al., 2009). Depending on the coding strand of a given gene, the transcription machinery either moves head-on (lagging strand genes) or co-directionally (leading strand genes) with respect to the movement of the replication machinery. The two types of encounters have different outcomes. In wild-type cells, high transcription of co-directional genes leads to replication stalling and restart (Merrikh et al., 2011). Furthermore, if cells lack RNA polymerase (RNAP) anti-backtracking factors, co-directional conflicts can lead to breaks in the DNA, at least on a plasmid in *Escherichia coli* (Dutta et al., 2011). However, in wild-type cells, many lines of evidence have demonstrated that when compared to co-directional conflicts, head-on conflicts are much more detrimental to both DNA replication and genomic stability (Boubakri et al., 2010; French, 1992; Liu and Alberts, 1995; Merrikh et al., 2015; Million-Weaver et al., 2015a, 2015b; Paul et al., 2013; Pomerantz and O’Donnell, 2008; Srivatsan et al., 2010). The reason for these drastically different negative consequences of the two types of conflicts is unclear.

Bacterial genomes tend to be organized such that most genes, especially high transcribed and essential genes, are expressed co-directionally with respect to DNA replication (Rocha and Danchin, 2003a, 2003b). This co-orientation bias has been attributed to both increased replication efficiency as well as decreased mutagenesis of genes when transcription occurs co-directionally with respect to replication (Paul et al., 2013). However, despite this strong co-orientation bias, which varies from species to species (Rocha and Danchin, 2003a, 2003b), many highly conserved (and some essential) genes remain in the head-on orientation (Paul et al., 2013). A number of these head-on genes play key roles in stress survival and are induced rapidly in response to environmental changes (Mostertz et al., 2004; Nicolas et al., 2012; Paul et al., 2013). Furthermore, it is clear that DNA replication continues upon induction of these genes since cells continue to grow after exposure to sub-lethal doses of many stressors, such as those that induce osmotic, oxidative, or cell wall stresses (Brill et al., 2011; Cao et al., 2005; Guariglia-Oropeza and Helmann, 2011; Mostertz et al., 2004; Nicolas et al., 2012). Additionally, though at a minimal level, replication stalling measurements have indicated that even in exponential phase, some (albeit mild) head-on conflicts occur (Merrikh et al., 2015; Million-Weaver et al., 2015a). Therefore, it is critical to identify mechanistically how head-on transcription disrupts replication and leads to genomic instability, and what strategies cells possess to resolve this potentially lethal problem. Yet, we know very little about gene orientation-specific mechanisms that disrupt DNA replication.

Given that during transcription and replication, movement of each machine along the template DNA causes over-winding of the two DNA strands ahead (Wu et al., 1988), when the two machines meet head-on, there should be excess positive supercoil formation between them (García-Muse and Aguilera, 2016; Mirkin and Mirkin, 2005). If so, this change in DNA topology and/or the stalling of transcription itself could either directly or indirectly generate hypernegatively supercoiled DNA behind RNAPs at the conflict region. It has been demonstrated that hypernegative supercoiling leads to the formation of RNA:DNA hybrids, referred to as R-loops when formed outside of the transcription bubble (Drolet et al., 1994; Masse and Drolet, 1999). These R-loops can lead to genomic instability and replication stalling (Gan et al., 2011; Lin and Pasero, 2012). Therefore, it is possible that instead of the classical model of RNAP itself directly being the culprit, a key factor contributing to the negative outcomes of head-on conflicts is the pervasive formation of R-loops at the conflict region.

R-loops are three-stranded structures consisting of a RNA:DNA hybrid and a displaced single-stranded DNA (Thomas et al., 1976), which can form upon transcription (Aguilera and García-Muse, 2012). In eukaryotes, many studies investigating R-loops have been carried out, and it is known that, in addition to effects on genomic instability and replication, these structures play important roles in gene regulation, mitochondrial DNA replication, and class-switch recombination (Aguilera and García-Muse, 2012; Costantino and Koshland, 2015; Santos-Pereira and Aguilera, 2015). In contrast, aside from potential roles in CRISPR immunity and replication restart outside of replication origins, little is known about R-loops with regards to genomic instability and replication in prokaryotes (Ivančić-Baće et al., 2012; Jore et al., 2011; Kogoma, 1997; Maduike et al., 2014; Rutkauskas et al., 2015). Furthermore, whether gene orientation (or replication-transcription conflicts) promotes R-loop formation or modulates the resulting effects of these structures in either eukaryotes or prokaryotes is unknown. Given that transcription is likely the most prevalent and disruptive obstacle to DNA replication, addressing this particular question is critical (Merrikh et al., 2015, 2011; Million-Weaver et al., 2015a).

The resolution of R-loops depends on the activity of highly conserved endonucleases known as RNases H, which cleave the RNA strand of RNA:DNA hybrids (Majorek et al., 2014; Tadokoro and Kanaya, 2009). If gene orientation impacts R-loop formation, RNases H may play important roles in the resolution of head-on replication-transcription conflicts. There are two major types of RNases H which are found across species: type 1 and type 2 (Ohtani et al., 1999a). Type 1 RNases H (including RNase HI) cleave the RNA strand of long RNA:DNA hybrids. Type 2 RNases H include RNase HII and HIII. Prokaryotic RNase HII removes single ribonucleotides that have been misincorporated into a DNA strand (Haruki et al., 2002). RNase HIII is categorized as a type 2 RNase H due to homology to RNase HII. However, the enzymatic activity of RNase HIII is more similar to the activity of RNase HI in that it cleaves the RNA strand of a long RNA:DNA hybrid (Ohtani et al., 1999b). In the Gram-positive model bacterium *Bacillus subtilis* there are functional RNase HII and HIII enzymes, encoded by the genes *rnhB* and *rnhC*, respectively (Itaya et al., 1999; Ohtani et al., 1999b). *B. subtilis* does not encode a functional RNase HI. However, it has been proposed that RNase HIII may serve the function of RNase HI in this organism (Itaya et al., 1999; Ohtani et al., 1999b). Whether RNases H play a role in resolution of head-on replication-transcription conflicts is unknown.

Here, we find that transcription in the head-on but not the co-directional orientation leads to pervasive RNA:DNA hybrid formation in *B. subtilis*. Our data indicate that *B. subtilis* RNase HIII is essential when cells are challenged with head-on conflicts. We find that pervasive R-loop formation blocks replication progression completely at the site of the conflict, specifically when transcription occurs head-on, but not co-directionally, with respect to DNA replication. This block in replication results in cell death. Additionally, without RNase HIII, we find that active DNA replication compromises the expression of specifically head-on, but not co-directionally oriented genes, further indicating that conflicts lead to pervasive R-loop formation at these regions. Importantly, we demonstrate that this mechanism plays a key role in stress survival. Most stresses strongly induce a large number of head-on genes, generating potentially lethal replication-transcription conflicts. Our data suggest that survival of these stresses, including oxidative, osmotic, and cell wall stress, requires the resolution of R-loops by RNase HIII at specifically head-on but not co-directionally oriented stress response genes.

## RESULTS

### R-loop formation is more prevalent when a gene is expressed in the head-on versus the co-directional orientation

We hypothesized that replication-transcription conflicts increase R-loop formation, specifically when they occur in the head-on orientation. To test this model, we measured the amount of RNA:DNA hybrids at an identical gene oriented either head-on or co-directionally with respect to DNA replication. We utilized engineered, site specific, inducible conflicts for these experiments. Briefly, the engineered conflict systems consist of reporter genes, placed under the control of inducible promoters, and inserted onto the chromosome of *B. subtilis*, in either the head-on or the co-directional orientation with respect to replication. These systems allow for the isolation of specifically gene orientation effects as the gene sequence, expression levels, and the chromosomal location where the conflict is induced are identical for the two constructs being compared in the experiments.

We built two different IPTG (Isopropyl β-D-1-thiogalactopyranoside) inducible reporter systems that either expressed the *lacZ* gene or the *luxABCDE* operon, P_*spank(hy)*_-*lacZ* and P_*spank(hy)*_- *luxABCDE*, respectively. We then introduced each of these constructs onto the chromosome at the *amyE* locus, in either the head-on or the co-directional orientation (Fig. 1A). To determine if R-loop formation is more prevalent in one or the other orientation, we measured the amount of RNA:DNA hybrids at these loci using RNA:DNA hybrid immunoprecipitations (DRIPs), followed by qPCRs. Briefly, we harvested cells containing the engineered constructs at mid-exponential phase, and generated lysates from these cells for DRIPs. We incubated the cell lysates with S9.6 antibody, which specifically binds RNA:DNA hybrids (García-Rubio et al., 2015). Upon elution, we quantified the amount of RNA:DNA hybrids in the DRIPs for either the head-on or the co-directionally oriented P_*spank(hy)*_-*lacZ* or P_*spank(hy)*_-*luxABCDE* constructs using qPCR. To measure the relative enrichment of hybrids at these engineered conflict regions, we normalized the signal for P_*spank(hy)*_-*lacZ* or P_*spank(hy)*_-*luxABCDE* to another locus (*yhaX*) on the chromosome. We found that more hybrids are formed when either *lacZ* or *luxABCDE* is expressed in the head-on compared the co-directional orientation (Fig. 1B and 1C). The relative DRIP signal for the head-on oriented genes was roughly two to three-fold higher compared to when they were oriented co-directionally with respect to replication (Fig. 1B and 1C). These results demonstrate that transcription in the head-on orientation increases the amount of RNA:DNA hybrid formation, strongly suggesting that head-on conflicts lead to pervasive R-loop formation.

**Figure 1.**
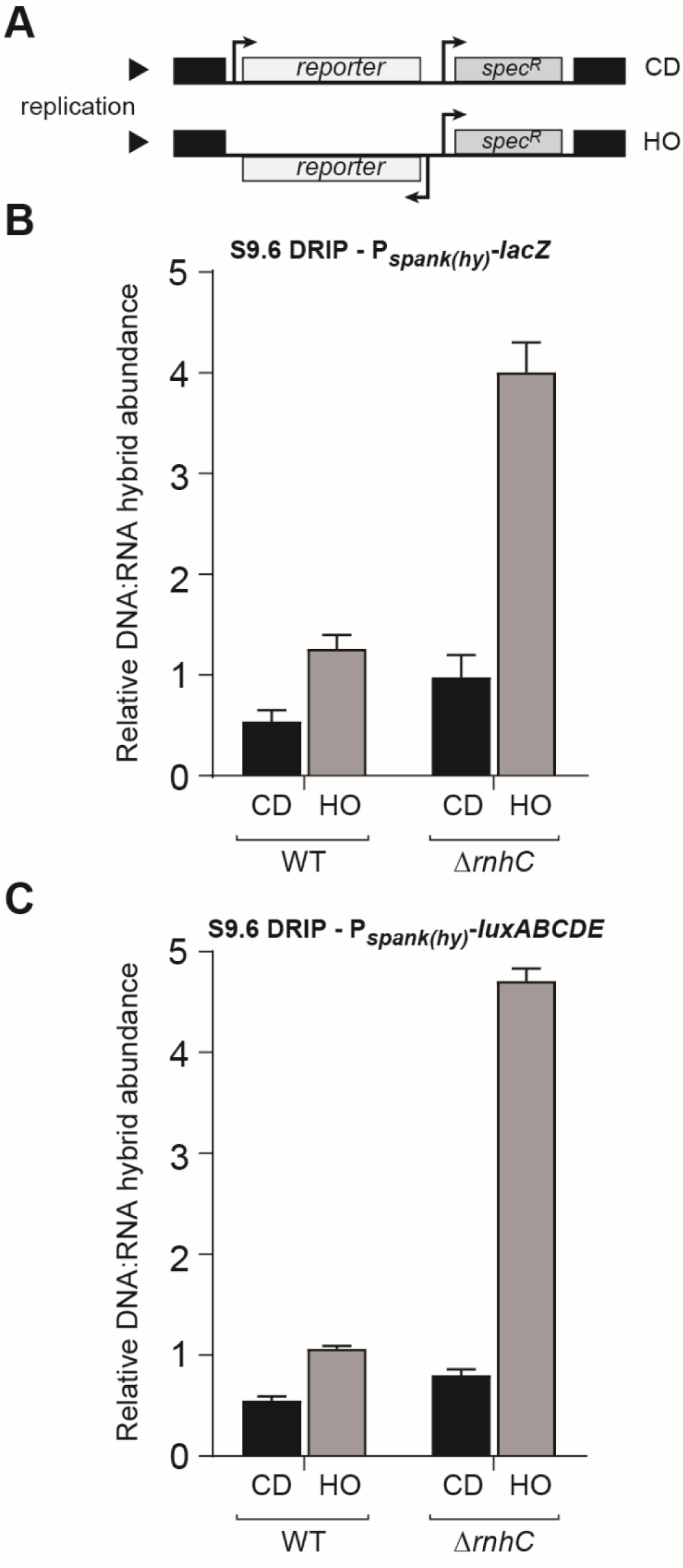
RNA:DNA Hybrids are prevalent at head-on genes. **(A)** Schematic of reporter gene integration in either the co-directional (CD) or head-on (HO) orientation with respect to replication. spec^R^, spectinomycin resistance gene. **(B)** qPCR of DNA from either wild-type cells (HM1300 (HO) or HM1416 (CD)) or cells lacking *rnhC* (HM2043 (HO) or HM2044 (CD)) pulled down with the S9.6 anti-RNA:DNA hybrid antibody. Genetic background is indicated below graph. Orientation of reporter gene *lacZ* is indicated directly below the bars: CD, Co-directional (black bars); HO, Head-on (gray bars). **(C)** qPCR of DNA from either wild-type cells (HM2195 (HO) or HM2197 (CD)) or cells lacking *rnhC* (HM2658 (HO) or HM2659 (CD)) pulled down with the S9.6 anti-RNA:DNA hybrid antibody. Genetic background is indicated below graph. Orientation of reporter gene *luxABCDE* is indicated directly below the bars: CD, Co-directional (black bars); HO, Head-on (gray bars).

### The absence of RNase HIII unveils pervasive R-loop formation at head-on genes

In *B. subtilis,* the gene *rnhC* codes for RNase HIII, which was shown to digest the RNA in long stretches of RNA:DNA hybrids or R-loops *in vitro* (Itaya et al., 1999; Ohtani et al., 1999b). If R-loops are processed *in vivo* by RNase HIII and the DRIP signal we detected was indeed a measurement of R-loops, in an *rnhC* knockout strain, the DRIP signal detected at the conflict region should increase. Furthermore, if head-on conflicts lead to pervasive R-loop formation, without *rnhC*, this increase in the DRIP signal should be more pronounced for the head-on oriented genes.

To address these questions, we used the DRIP assay to measure the abundance of RNA:DNA hybrids in the two orientations, in cells lacking *rnhC*. We found that without *rnhC*, more RNA:DNA hybrids form at the region harboring either the *lacZ* or the *luxABCDE* constructs, in both orientations (Fig. 1B and 1C). However, this increase in the DRIP signal was significantly more pronounced for the head-on orientation (about four-fold) compared to the co-directional orientation (about two-fold) (Fig. 1B and 1C). This finding suggests that expression in the head-on orientation promotes R-loop formation, and that RNase HIII activity is needed to resolve these structures.

### Pervasive R-loop formation at head-on genes can completely block replication fork progression and prevent replication restart

Given that we found an increased enrichment of RNA:DNA hybrids at genes expressed head-on to replication, we wondered if R-loops exacerbate head-on conflicts by blocking DNA replication specifically in this orientation. To address this model, we measured replication stalling in the presence or absence of *rnhC* via: 1) chromatin immunoprecipitations (ChIPs) of the replicative helicase DnaC, 2) 2D gel electrophoresis assays (2D gels), and 3) marker frequency assays (DNA copy measurement through deep sequencing).

In the ChIP experiments, the relative association of the replicative helicase (DnaC) with the conflict regions compared to control loci was measured as previously described (Merrikh et al., 2011; Million-Weaver et al., 2015a). Replisome proteins, such as DnaC, are not expected to preferentially associate with any given locus unless there is replication slow-down due to an obstacle (this holds true in an asynchronous population of cells). Thus, increased relative association of DnaC with a particular chromosomal locus is most likely indicative of replisome stalling or restart at that region (Merrikh et al., 2011). Similar to our prior findings, we detected a roughly 2-3 fold increase in association of DnaC with the head-on compared to the co-directionally oriented P_*spank(hy)*_-*lacZ* reporter gene (Fig. 2A). In the absence of *rnhC*, there was a significant increase in DnaC association only with the head-on, and not the co-directionally oriented P_*spank(hy)*_-*lacZ* (Fig. 2A). This indicates that R-loop formation is specifically problematic for DNA replication at head-on transcribed regions.

**Figure 2.**
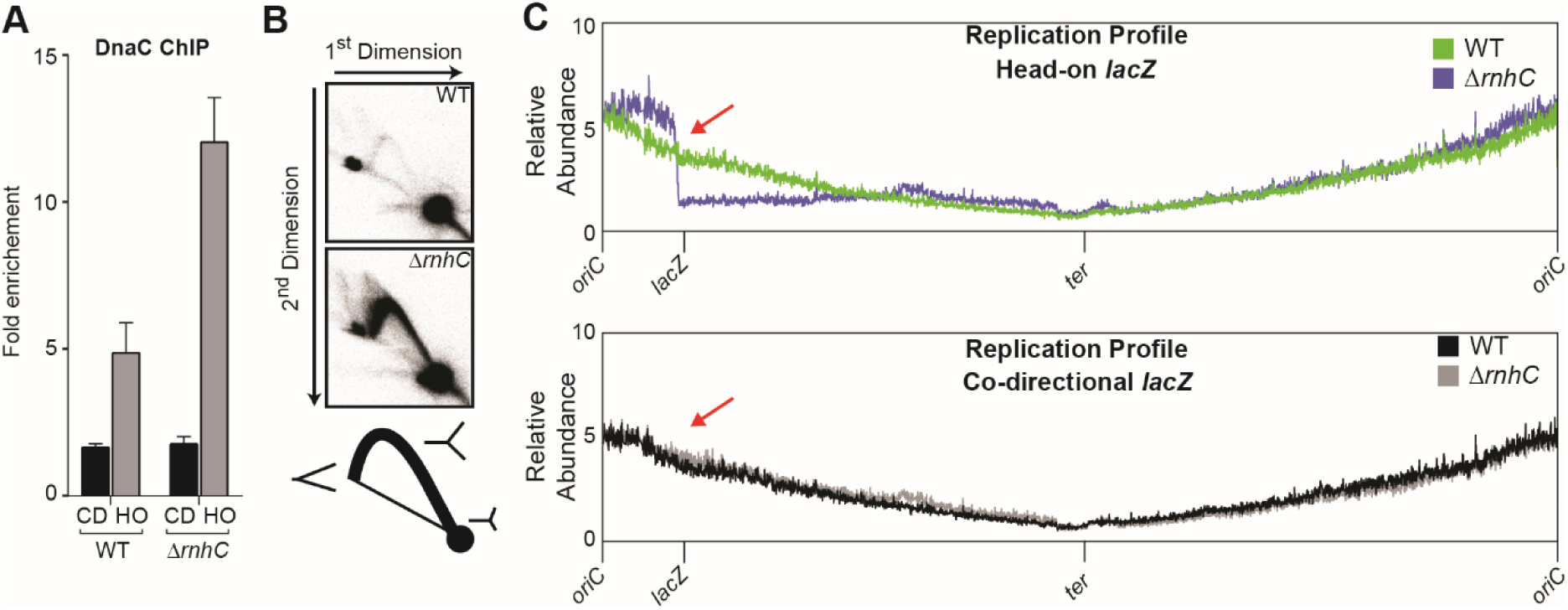
Stable R-loops that form at head-on genes block replication. **(A)** qPCR analysis, targeting the reporter gene *lacZ,* of either WT (HM1300 (HO) or HM1416 (CD)) or Δ*rnhC* (HM2043 (HO) or HM2044 (CD)) cells using polyclonal antibodies against the replicative helicase DnaC. **(B)** 2-dimensional gel electrophoresis analysis of the EcoRV digested fragment containing the head-on reporter gene *lacZ* and downstream region. Top panel is DNA isolated from wild-type cells (HM1300). Middle panel is Δ*rnhC* (HM2043). Bottom panel contains a cartoon schematic of where various branched structures should run on the gel. **(C)** Deep sequencing analysis of genomic DNA isolated from either WT or Δ*rnhC* cells containing the reporter gene, *lacZ* (indicated by the red arrow), in either the Head-on (top panel, HM1300 (wild-type); HM2043 (Δ*rnhC)*) or Co-directional (bottom panel, HM1416 (wild-type); HM2044 (Δ*rnhC)*) orientation. The x-axis indicates chromosomal locations and the y-axis is the abundance of reads relative to the total number of reads in the sequencing library.

We obtained similar results when we assessed replication stalling using 2D gels, which allow for direct analysis of replication intermediates trapped at a given chromosomal locus (Friedman and Brewer, 1995). In these experiments, cells are gently lysed in agarose plugs to preserve DNA structures and specific regions are isolated using restriction digests. Specific chromosomal locations are detected using radio-labeled ssDNA probes. Branched replication fork structures are detected as Y-arcs on 2D gels (Fig. 2B). As we previously reported, replication intermediates only appear within the engineered conflict regions at high transcription levels, and in particular in the head-on orientation (Merrikh et al., 2015). Consistent with the DnaC ChIP results, without *rnhC*, the Y-arc intensity for the head-on oriented P_*spank(hy)*_-*lacZ* region increases significantly (Fig. 2B), which demonstrates that replication fork progression through head-on transcribed regions is blocked due to R-loop formation.

To determine how severe the observed stalling is, and whether replication progresses beyond the conflict region, we performed marker frequency assays. We harvested cells in mid-exponential phase after inducing P_*spank(hy)*_-*lacZ* with IPTG, and extracted genomic DNA. We used deep sequencing to quantify and map the relative abundance of all genomic loci in these samples. The frequency of mapped reads can be used as a proxy for DNA copy number or replisome progression. Strikingly, we found that, without RNase HIII, there is a sharp drop in DNA copy number beginning at the head-on conflict region, which lasts throughout the remainder of that arm of the chromosome (Fig. 2C). In contrast, there was no drop in marker frequency without *rnhC* when the *lacZ* gene was transcribed in the co-directional orientation. We note that there was an increased signal surrounding the origin in the *rnhC* knockout strain containing head-on (but not co-directional) *lacZ*, which could be indicative of increased replication initiation. Additionally, we found a small increase in DNA copy number at the region encoding for phage SPβ without *rnhC*, but this increase was independent of reporter orientation. The orientation independence of this effect suggests that the increased copy number of phage DNA is a general response to lacking RNase HIII. Our results indicate that if not resolved, R-loop formation at, specifically, head-on oriented genes can completely block replication and subsequent replication restart.

### Pervasive R-loop formation caused by head-on replication-transcription conflicts is lethal

Given that we detected a complete replication stop at the head-on but not the co-directionally oriented conflict regions, we predicted that head-on conflicts would be lethal in the absence of *rnhC*. To test this model, we examined the impact of knocking out *rnhC* on survival of cells containing the head-on versus the co-directionally oriented P_*spank(hy)*_-*lacZ* and P_*spank(hy)*_-*luxABCDE* constructs. Briefly, cells were grown to mid-exponential phase, serially diluted, and then spotted onto plates that either did or did not contain the inducer of the P_*spank(hy)*_ promoter, IPTG. *rnhC* knockouts have a relatively mild growth defect as the colony size for these mutants is generally smaller than that for wild-type (Fig. 3). We found that, in the absence of induction (no IPTG), cells containing the reporters in either orientation did not have any additional growth defects, regardless of whether the strain did or did not contain *rnhC* (Fig. 3A and 3B). Furthermore, induction of either the *lacZ* gene or the *luxABCDE* operon by IPTG did not produce any additional survival defects in wild-type cells. Remarkably, however, without *rnhC*, cells containing the head-on but not the co-directionally oriented reporters showed severe growth defects that were dependent on expression levels (controlled by various amounts of IPTG addition) (Fig. 3A and 3B). At full induction (1mM IPTG), we were unable to detect any growth for cells containing specifically the head-on but not the co-directionally oriented P_*spank(hy)*_-*lacZ* or P_*spank(hy)*_-*luxABCDE* construct (Fig. 3A and 3B). The reproducibility of these results with two completely different genes controls for potential artifacts of spurious protein production from these reporter genes. The essentiality of RNase HIII depends on both orientation and transcription, strongly suggesting that head-on replication-transcription conflicts lead to pervasive R-loop formation that as we determined, completely blocks DNA replication. The observed lethality of expressing a single gene in the head-on orientation is consistent with the stalling measurements.

**Figure 3.**
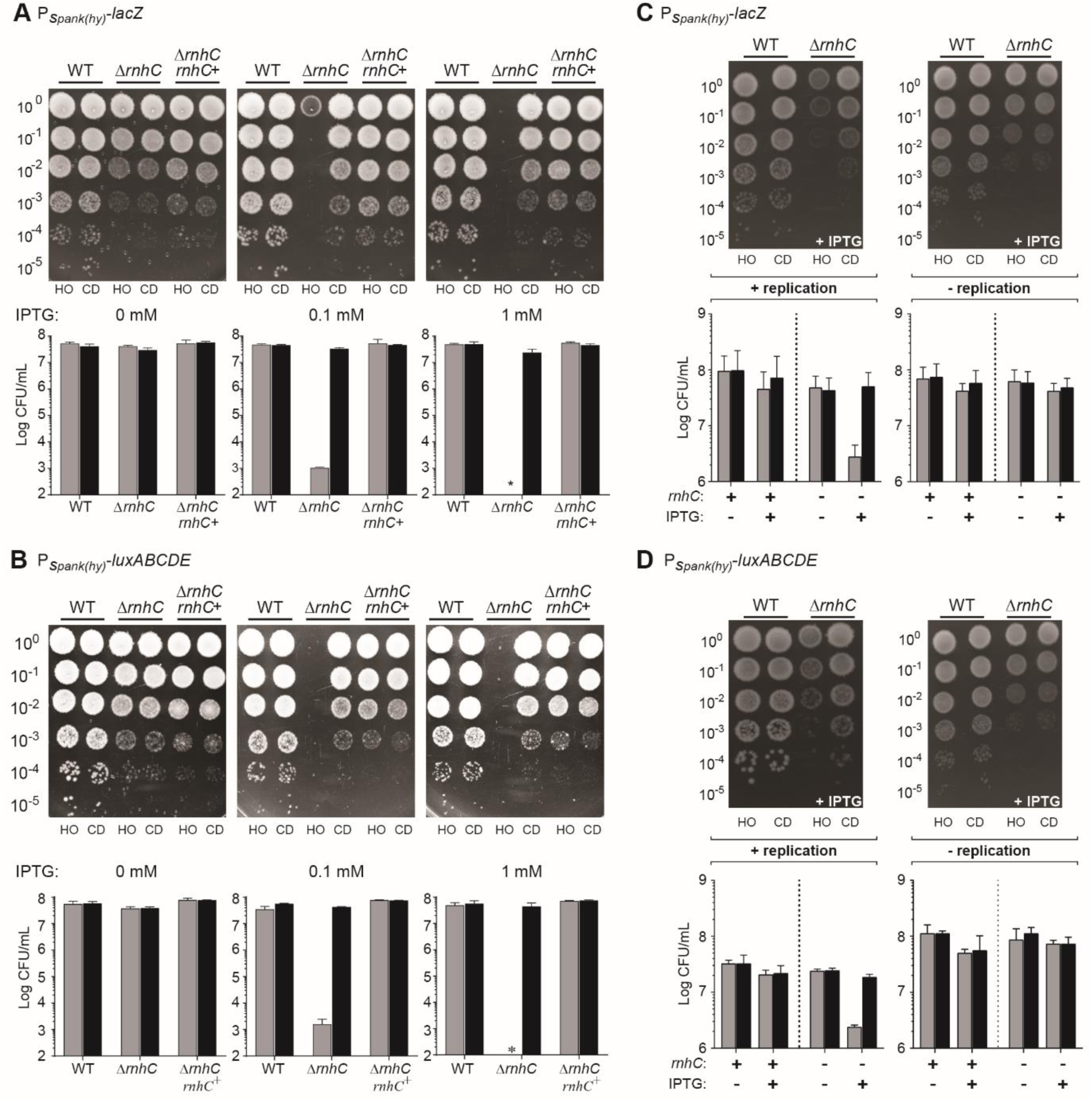
Unresolved R-loops at head-on genes cause cell death. **(A)** Top panel, representative plates of survival assays for either wild-type (WT) cells (HM1300, HM1416), or Δ*rnhC* cells (HM2043, HM2044), or Δ*rnhC* cells with *rnhC* expressed at an ectopic locus (Δ*rnhC rnhC^+^*, HM2656, HM2657) harboring the IPTG-inducible reporter gene *lacZ* in either the HO or CD orientation plated on LB plates contain 0, 0.1 mM or 1 mM IPTG. Bottom panel contains the quantification of at least 4 biological replicates. **(B)** Top panel, representative plates of survival assays for eitherWT cells (HM2195, HM2197), or Δ*rnhC* cells (HM2658, 2659), or *rnhC* deletion with *rnhC* expressed at an ectopic locus (Δ*rnhC rnhC^+^*, HM2706, HM2707) harboring the IPTGinducible reporter gene *luxABCDE* in either the HO or CD orientation plated on LB plates contain 0, 0.1 mM or 1 mM IPTG. Bottom panel contains the quantification of at least 4 biological replicates. **(C)** Top panel, representative plates of survival assays for either WT cells (HM1300, HM1416), or Δ*rnhC* cells (HM2043, HM2044) harboring the IPTG-inducible reporter gene *lacZ*. **(D)** Top panel, representative plates of survival assays for either WT cells (HM2195, 2197), or Δ*rnhC* cells (HM2658, 2659) harboring the IPTG-inducible reporter gene *luxABCDE*.

If R-loops form as a result of head-on collisions between the replication and transcription machineries, then presumably, if a gene is transcribed in the absence of replication, R-loop formation will not be toxic. We tested this hypothesis by expressing our engineered reporter constructs (*lacZ* and *luxABCDE*) in the two orientations, either during active replication (exponentially growing cells) or in the absence of replication (cells in stationary phase). We transiently induced the reporters with IPTG, and then spotted the cells onto plates lacking IPTG. When this transient expression experiment was performed in actively growing wild-type cells, we did not detect any decrease in viability regardless of whether the genes were expressed in the head-on or the co-directional orientation (Fig. 3C and 3D). However, in cells lacking RNase HIII (*rnhC* mutants), there was a roughly one to two logs decrease in CFUs/mL, but only when transcription was induced in actively replicating cells and only when the reporter was expressed in the head-on orientation (Fig. 3C and 3D). In contrast, this orientation-specific viability defect was not observed when the genes were induced in non-replicating cells, strongly suggesting that active replication is required for the orientation-specific R-loop toxicity (Fig. 3C and 3D). Based on these results, the lethality of head-on transcription most likely stems from pervasive R-loop formation caused by head-on replication-transcription conflicts.

### R-loop formation at head-on but not co-directional genes leads to gene expression defects

R-loops have been associated with impaired RNAP movement and gene expression defects (Santos-Pereira and Aguilera, 2015; Skourti-Stathaki and Proudfoot, 2014). Given that we found R-loops are more prevalent and/or stable in the head-on orientation, we anticipated that head-on, but not co-directional gene expression, would suffer without RNase HIII. To test this hypothesis, we measured the abundance of *lacZ* or *luxABCDE* mRNA using RT-qPCR in cells with and without RNase HIII. When *rnhC* was absent, expression of the head-on oriented genes decreased significantly (Fig. 4A and 4B). When the same genes were expressed in the co-directional orientation, there was no difference in their mRNA abundance. Furthermore, the decrease in mRNA levels of the head-on reporter genes depended on active DNA replication: when we repeated these experiments in non-growing cells (stationary phase cultures), the head-on genes were induced significantly with IPTG addition, however, the observed gene expression defect without *rnhC* was no longer detectable (Fig. 4C). These data are consistent with the results presented above, strongly suggesting that R-loop formation is specifically problematic in the head-on orientation due to encounters between the replication and transcription machineries.

**Figure 4.**
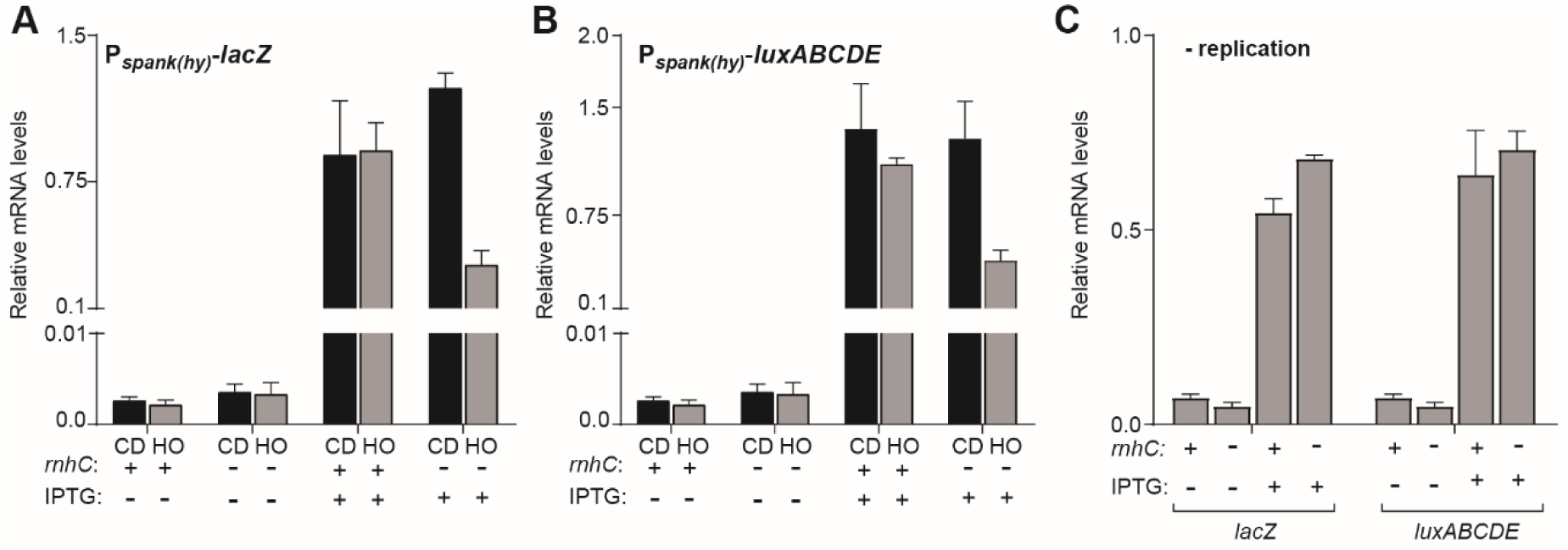
R-loops disrupt head-on gene expression. qPCR analysis of reporter genes in either WT (*lacZ*: HM1300, HM1416; *luxABCDE*: HM2195, 2197) or Δ*rnhC* (*lacZ*: HM2043, HM2044; *luxABCDE*: HM2658, HM2659) cells. The y-axis indicates the abundance of reporter gene mRNA relative to the housekeeping gene *dnaK.* Black bars indicate cells harboring the reporter gene in the CD orientation, gray bars indicate cells harboring the reporter gene in the HO orientation. **(A-B).** Cells were induced for 1 hour with 1 mM IPTG and harvested during log phase growth. **(C).** Cells were induced for 1 hour with IPTG (1 mM per 0.5 OD600 units) and harvested in stationary phase.

### Efficient expression of head-on stress response genes requires R-loop resolution

Many stress response genes are coded in the head-on orientation in bacteria (Table 1 and (Mostertz et al., 2004; Nicolas et al., 2012; Paul et al., 2013)). Given our findings that R-loop formation at a single, highly expressed gene can lead to significant gene expression defects, we postulated that resolution of R-loops at head-on conflict regions will be critical for proper expression of head-on oriented stress response genes. We predicted that upon exposure to stress, the expression of these head-on genes will generate a significant number of conflicts and pervasive R-loop formation that if not resolved by RNase HIII, will compromise expression of these key stress response genes. To test this model, we selected representative genes known to be highly induced upon exposure to stress in growing cells, in both orientations, and measured their expression levels during three different stress conditions: lysozyme (cell wall stress), paraquat (oxidative stress), or salt (osmotic stress). Each of these stresses induces an array of genes (Mostertz et al., 2004; Nicolas et al., 2012). We analyzed both head-on (lysozyme: *sigV* and *dltA*, paraquat: *katA* and *aphC*, salt: *proH* and *ysnF*) and co-directional genes (lysozyme: *ddl* and *murF*, paraquat: *ykuP* and *dhbB*, salt: *katE* and *yjgC*). We found that in cells lacking *rnhC*, there is a defect in expression of only the head-on oriented stress response genes we analyzed, and only when cells were actively replicating (Fig. 5). In contrast, in non-growing cells, there was no difference in expression levels of the head-on stress response genes when comparing wild-type cells to those lacking *rnhC*. When we exposed cells to these stresses under conditions of growth-arrest, the stress response genes were induced as expected, indicating that the stress treatment was effective. Additionally, we did not observe an expression defect for any of the co-directionally oriented genes without *rnhC* (Fig. 5). Note that we only analyzed a few genes under each condition, thus these results need not apply to all genes. However, the results of these experiments are consistent with the reporter experiments presented above, suggesting that pervasive R-loop formation is a general principle of gene orientation that is independent of genomic loci, sequence context, promoter nature, or the degree of induction. Given the results of the stress exposure experiments, a robust stress response in bacteria most likely relies on efficient removal of R-loops generated at head-on genes due to collisions with replication.

**Figure 5.**
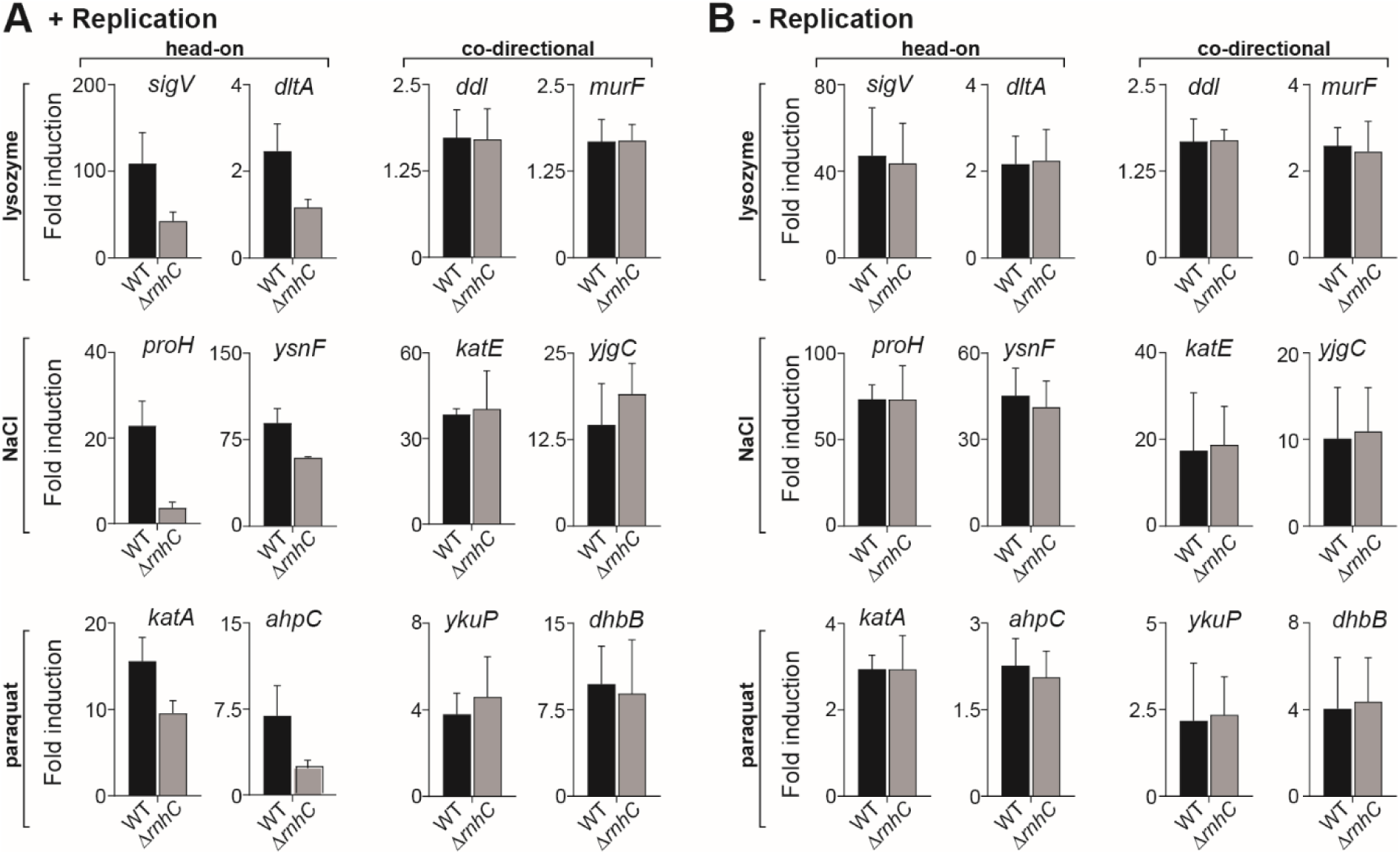
R-loops block expression of head-on stress response genes. qPCR analysis of reporter genes in either WT (HM1) or Δ*rnhC* (HM2655) cells. The y-axis indicates the fold induction of the indicated gene’s mRNA relative to the housekeeping gene *dnaK* after the indicated treatment compared to untreated cultures. Black bars indicate wild-type cells and gray bars indicate Δ*rnhC* cells. **(A)** Cells were treated and harvested during exponential growth. **(B)** Saturated, stationary phase cultures were treated and harvested.

**Table 1.**
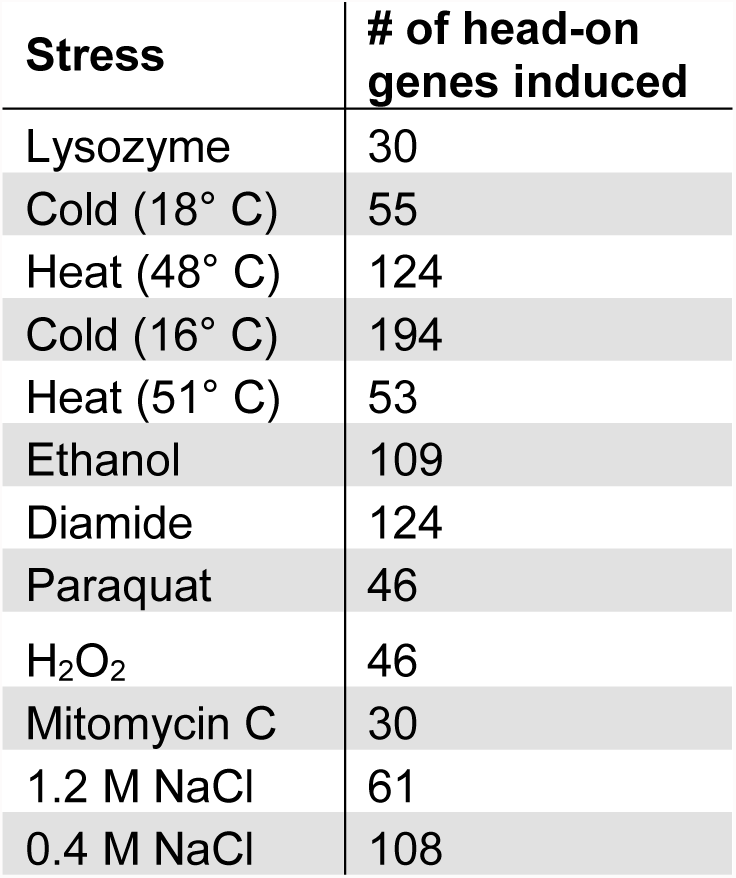
Response to stress induces expression of head-on genes. Data were compiled from (Guariglia-Oropeza and Helmann, 2011; Mostertz et al., 2004; Nicolas et al., 2012).

### Efficient survival of stress requires resolution of R-loops at head-on conflict regions

It is possible that resolution of head-on replication-transcription conflicts by R-loop processing is needed for stress survival, either because of the need for efficient expression of head-on stress response genes or because expression of these head-on genes results in replication stalling. To test this model, we chronically exposed cells to physiologically and environmentally relevant stresses including lysozyme, oxidative stress, and osmotic shock by performing viability assays on agar plates supplemented with each of these various stressors. We compared wild-type cells to those lacking *rnhC* and found that indeed, efficient survival of all these stresses requires *rnhC* (Fig. 6A). Without *rnhC*, chronic exposure to different stresses decreased viability as measured by CFUs/mL by one to three orders of magnitude (Fig. 6A).

**Figure 6.**
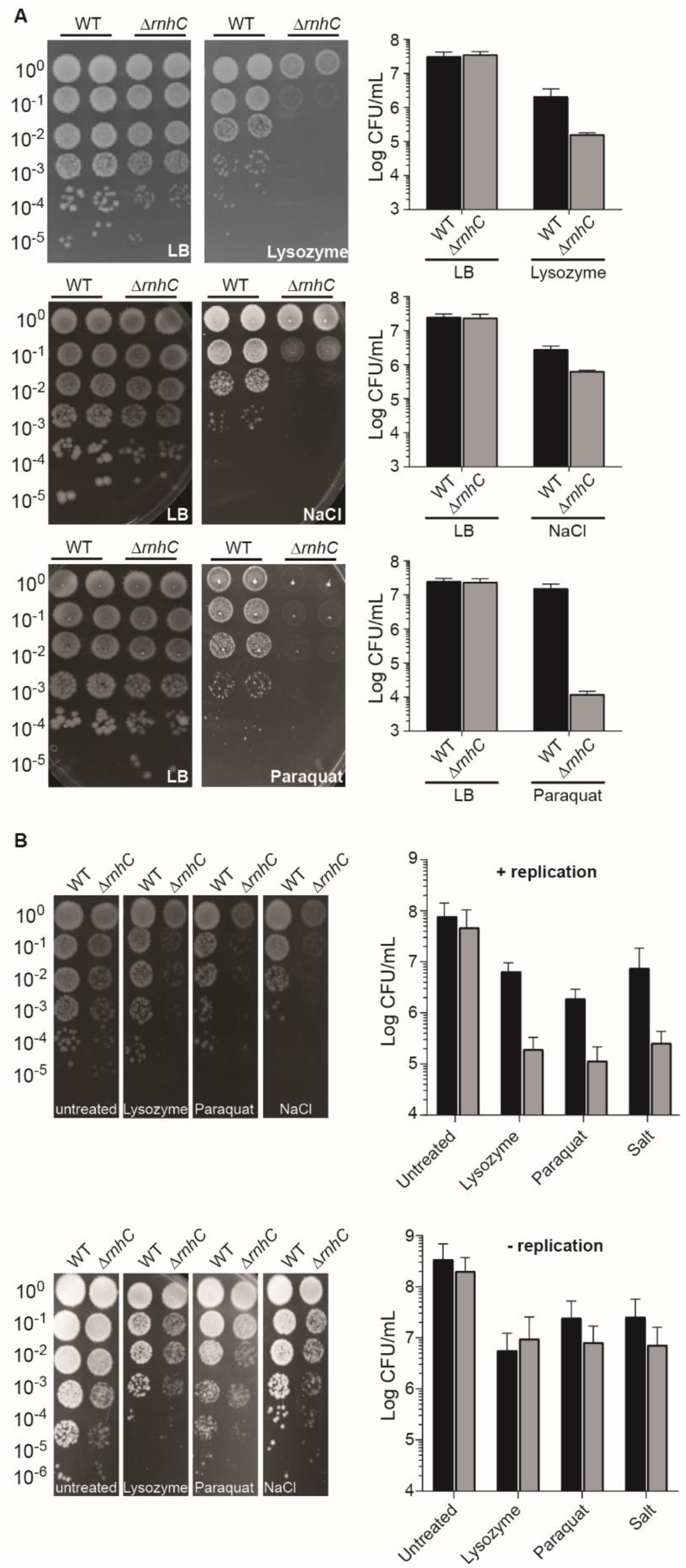
Resolution of R-loops at head-on genes is required for survival of stress. **(A)** Representative plates (left) and quantification (right) from chronic survival assays of either WT (HM1) or Δ*rnhC* (HM2655) cells plated on LB or LB containing 50 μg/mL of lysozyme (top), 700 mM NaCl (middle) or 70 μM paraquat (bottom). **(B)** Representative plates (right) and quantification of acutely treated survival assays. Top panel are results from cells treated in exponential phase. Bottom panel are results obtained from cells treated in saturated stationary phase cultures. Type of treatment is indicated on bottom of plate or under the x-axis. Black bars on the quantification plots represent wild-type cells and gray bars indicate Δ*rnhC* cells.

We tested whether the cell survival defect observed for the *rnhC* mutants depends on active replication. We acutely treated cells with the various stressors, either when they were actively replicating, in exponential phase, or not replicating, in stationary phase. We then measured cell viability by enumerating CFUs/mL as described above. When cells were treated with stresses during either active growth or non-growth, in stationary phase, there was a significant (roughly 90% - 99% decrease in CFUs/mL) survival defect in both wild-type and *rnhC* knockout cells, indicating that the acute stress treatments were effective. When actively replicating cells were treated with stresses, cells lacking *rnhC* were more sensitive to the corresponding stress (Fig. 6B). However, when we performed the acute stress treatments in non-replicating cells, we no longer observed a difference in survival of wild-type versus *rnhC* mutants. Taken together with the gene expression data, the results of these survival assays strongly suggest that resolution of R-loops formed as a result of head-on replication-transcription conflicts is essential for efficient survival of many different stresses.

## DISCUSSION

Our data suggest that gene orientation strongly impacts R-loop formation due to conflicts between transcription and replication. If not resolved by RNase HIII, R-loops can completely stall replication at head-on genes, resulting in cell death. Additionally, these R-loops block efficient expression of head-on genes. Based on our results, we propose a model where R-loops form during collisions between the replication and transcription machineries. As the replisome approaches a head-on transcription unit, RNAPs begin to stall, either due to increased positive supercoiling or a direct collision with the replisome. Nascent mRNAs from the stalled RNAPs might then invade the DNA duplex efficiently, which would presumably be open due to active transcription (and potentially hypernegative supercoiling). Once formed, these RNA:DNA hybrids would require RNase HIII activity for resolution. If left unresolved, these stable RNA:DNA hybrids block replication either directly or by stabilizing RNAPs, which could then act as a barrier to replication. RNA:DNA hybrids are extremely stable, and therefore, excess R-loop formation caused by conflicts could have long-term consequences that extend beyond the time of fork passage through the transcribed region. We cannot rule out that our DRIP signal stems from unprocessed Okazaki fragments at the site of the head-on conflict. However, because *rnhC* is not essential, cells likely have other means of processing Okazaki fragments, making this scenario unfavorable. However, *rnhC* mutants form smaller colonies, which could be a result of inefficient Okazaki fragment processing. Alternatively, this defect in colony morphology may be due to head-on replication-transcription conflicts that occur during “stress-free” or exponential growth. Given our findings, we prefer the latter model.

The question of why head-on conflicts are more detrimental than co-directional conflicts has been long standing. Our data strongly suggest that a key difference between head-on and co-directional replication-transcription conflicts is the local alteration in nucleic acid structures, mainly, pervasive R-loop formation. There are physical differences between what occurs at replication-transcription conflict regions when a gene is encoded on the lagging strand (head-on orientation) compared to the leading strand (co-directional orientation). For example, the topology of the DNA at the conflict region has been predicted to be different in head-on compared to co-directional collisions (García-Muse and Aguilera, 2016; Mirkin and Mirkin, 2005). The “Twin-Domain” model of DNA topology predicts that positive supercoiling is generated in front of both RNAP and the replication machinery (Wu et al., 1988). As these two complexes move toward one another, positive supercoiling would be expected to transiently build up at the site of the conflict. This build up presumably would not be present at the regions of co-directional encounters.

Our results show that *in vivo*, the replication fork can be completely blocked by stable RNA:DNA hybrids and/or RNAPs stabilized by hybrids. Given that the replicative helicase encircles the lagging strand, it may not be able to translocate across R-loops formed at head-on genes. Alternatively, R-loops could stabilize RNAPs in such a way that they cannot be cleared and consequently act as barriers to the replisome. Remarkably, cells lacking RNase HIII apparently cannot restart replication. A small increase in DNA copy number far downstream of the head-on conflict region is observed in the replication profiles presented above, which could represent a small subpopulation of cells undergoing R-loop dependent replication restart near the terminus. One caveat is that this signal increase coincides with the location of a lysogenized phage and could also represent a lytic phage cycle initiated by host replication arrest. The lack of replication restart upon R-loop formation at conflict regions could stem from unavailability of the necessary DNA for restart proteins to bind. The recruitment and binding of the key replication restart protein PriA requires either a D-loop (generated by the recombination protein RecA) or a stalled fork structure (Liu and Marians, 1999). Perhaps R-loops mask these DNA structures by making one of the two DNA strands unavailable to restart factors. R-loops have been proposed to initiate replication restart outside the origin (Boubakri et al., 2010; Maduike et al., 2014). However, our data demonstrate that R-loop-mediated restart cannot occur from head-on genes, at least in *B. subtilis*.

Given that all organisms contain head-on genes, experience conflicts, and possess RNase H enzymes, R-loops are likely to be a universal problem across species. We find that there is a correlation between the existence of specifically RNase HIII and gene orientation bias: bacteria with a high bias toward co-orientation of genes, such as *B. subtilis*, contain RNase HIII (Figure 7). This suggests that perhaps in organisms without such a strong bias, such as *E. coli*, R-loops are not as detrimental, or that redundant R-loop processing mechanisms exist. Indeed, in *E. coli*, rDNA inversions are not as disruptive to growth as in *B. subtilis* (Boubakri et al., 2010; Zhang et al., 2014). However, R-loops must be problematic to some degree even in low bias organisms such as *E. coli* given that modulating RNase HI levels can impact survival defects associated with rDNA inversions (Boubakri et al., 2010; Zhang et al., 2014). Additionally, *E. coli* encodes accessory helicases, which may act as redundant mechanisms to remove R-loops, again suggesting that these structures are indeed problematic across species (Boubakri et al., 2010). These redundant mechanisms are also conserved: we have previously shown that in *B. subtilis*, the accessory helicase PcrA is needed for replication through transcription units (Merrikh et al., 2015). The homolog of PcrA in *E. coli,* UvrD, has been shown to unwind RNA:DNA hybrids *in vitro* (Matson, 1989). In yeast, the PcrA homolog Pif1 has been shown to preferentially unwind RNA:DNA hybrids over DNA:DNA hybrids (Boulé and Zakian, 2007; Chib et al., 2016). Interestingly however, despite the established role of helicases in R-loop processing, here we find that helicases are not sufficient to process excess R-loop formation, specifically in head-on genes. Perhaps, this is the reason for the presence of RNase HIII in organisms with higher co-orientation biases.

**Figure 7.**
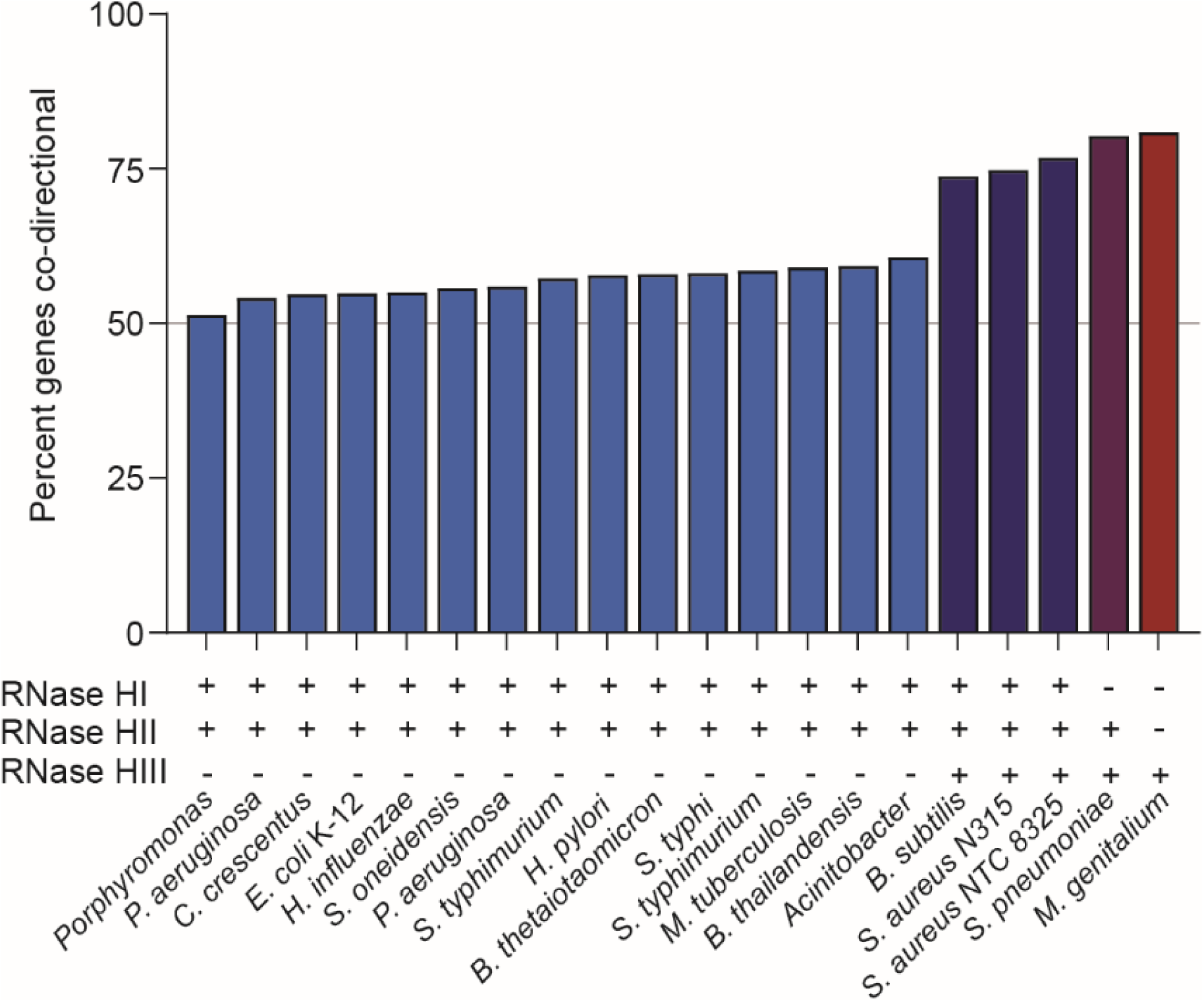
Co-orientation of genomes correlates to presence of RNase HIII. Data were compiled from (Zheng et al., 2015) and (Kochiwa et al., 2007). The y-axis represents the percent of genes on the chromosome that are co-oriented with respect to replication. Different strains and the presence (+) or absence (-) of genes encoding the different RNase H enzymes are indicated along the x-axis.

In this work, we have demonstrated that replication-dependent R-loop formation can inhibit efficient expression of head-on genes, which is needed for survival of common stresses. Thus, head-on conflicts, and the resulting over-abundance of R-loops must be a life threatening problem in nature. An imminent inactivating mutation in *rnhC* could be lethal during stress exposure. These findings resurrect a question we posed previously: if expression of head-on genes has the potential to completely block replication via R-loop formation, and cells need elaborate mechanisms to survive these problems, why are there genes encoded on the lagging strand at all? This paradox can be explained in light of evolution. In our previous work, we demonstrated that genes in the head-on orientation accumulate mutations faster over evolutionary time scales (Paul et al., 2013). Therefore, the conservation of the head-on orientation for some genes could potentially serve as a source for increased genetic diversity. This could potentially generate the genetic diversity needed for a population to survive unforeseen stresses.

R-loops seem to lead to severe problems when they form in genes that are transcribed opposite to replication. In turn, RNase H enzymes, which are highly conserved from bacteria to humans, become essential for resolution of life-threatening head-on conflicts between replication and transcription. Given that bacteria face stresses that induce head-on genes routinely in the environment, including during times of growth such as host invasion and *in vivo* replication within eukaryotes, we predict that head-on conflict resolution is a key process that supports bacterial life. In particular, resolution of R-loop formation in response to various stresses is likely required for bacteria to sustain life in nature. R-loop formation at head-on genes therefore is a dangerous problem, and although not always identified as “essential” in the laboratory, RNase H enzymes and their homologs are most likely required for life under the diverse environmental settings in the wild. Given the conservation of RNase H enzymes and the universal nature of replication-transcription conflicts, the basic principles of what we have discovered should also apply to eukaryotic organisms.

## MATERIALS AND METHODS

### Strains and growth conditions

All *B. subtilis* strains were constructed in the HM1 (JH642) (Brehm et al., 1973) *Bacillus subtilis* background. The *rnhC::mls* mutant (HM711) was obtained from the *Bacillus* genetic stock center (Columbus, OH). To move the *rnhC::mls* allele, genomic DNA was extracted from HM711 using a commercially available kit (Thermo) and used to transformed into HM1 (and its derivatives with reporter constructs) as per standard protocol (Cutting and Harwood, 1990). Strains were streaked on LB agar plates and supplemented with antibiotics where appropriate. Precultures were inoculated from single colonies into 2 or 5 mL of LB broth and incubated at 37° C with shaking (260 RPM). Precultures were used to inoculate experimental cultures which were grown and treated as indicated for each different experiment in the Method Details section.

*E. coli* DH5α was used to propagate recombinant DNA vectors. Transformations were done using heat shock of competent *E. coli*. *E. coli* cultures were grown at 37° C with shaking (260 RPM) in LB supplemented with 50 μg/mL carbenecillin where appropriate. All plasmid vectors were purified using a commercially available plasmid extraction kit (Thermo).

### Strain constructions

#### pHM170

The *lacZ* gene from pKG1 (Million-Weaver et al., 2015a) was amplified by PCR using primers HM1463 and HM1464. This PCR amplicon was digested with HindIII and SphI (Thermo) restriction enzymes and ligated into HindIII and SphI digested pDR111. The ligation reaction was subsequently transformed into *E. coli* DH5α and the construct was confirmed by PCR and DNA sequencing (Genewiz). The plasmid was subsequently purified and transformed into competent HM1 to generate strain HM1300.

#### pHM181

The *lacZ* gene from pHM170 was amplified by PCR with primers HM2120 and HM2121. The PCR amplicon was digested with SphI and EcoRI and ligated into SphI and EcoRI digested pDR111. The ligation reaction was subsequently transformed into *E. coli* DH5α and the construct was confirmed by PCR and DNA sequencing (Genewiz). The plasmid was subsequently purified and transformed into competent HM1 to generate strain HM1416.

#### pHM321

The *lux* operon was amplified by PCR from pAH60 (Ferguson et al., 2007) using primers HM2747 and HM2748. The PCR amplicon was digested with EagI and NheI and ligated into pDR111 that was digested with EagI and NheI. The ligation reaction was subsequently transformed into *E. coli* DH5α and the construct was confirmed by PCR and DNA sequencing (Genewiz). The plasmid was subsequently purified and transformed into competent HM1 to generate strain HM2197.

#### pHM271

The *lacA*::P*_rnhC_-rnhC* complementation construct was built by amplification of the native *rnhC* locus, including the native *rnhC* promoter (P_*rnhC*_) and terminator, with primers HM2406/2407, tagged with EcoRI and XhoI restriction sites. The *rnhC* PCR amplicon was digested with EcoRI and XhoI (Thermo) and ligated into the EcoRI and XhoI sites of pMMB752. The resulting vector, pHM271 (*lacA*::P*_rnhC_-rnhC*) was subsequently transformed into *E. coli* DH5α and confirmed by PCR before transformation into competent *B. subtilis.* Integration at *lacA* was confirmed by colony PCR.

### Viability assays - chronic treatments

Strains were struck on LB plates supplemented with the appropriate antibiotic from freezer stocks and incubated overnight at 37° C. Single colonies were used to inoculate 2 mL LB cultures in glass tubes. The cultures were grown at 37° C with shaking (260 RPM) to OD600 = 0.5-1.0. Precultures were adjusted to OD 0.3 and then serially diluted in 1x Spizzen’s Salts (15 mM ammonium sulfate, 80 mM dibasic potassium phosphate, 44 mM monobasic potassium phosphate, 3.4 mM trisodium citrate, and 0.8 mM magnesium sulfate). 5ul of each dilution was plated onto LB plates and incubated at 30° C overnight. For survival assays with reporter strains, LB plates were either supplemented or not with various concentrations of IPTG (1mM, 0.1mM) For viability assays of cells chronically treated with stressors, plates were supplemented with various concentrations of lysozyme (50ug/mL), salt (700 mM), and paraquat (70 mM). Plates were imaged with a BioRad Gel Doc™ XR+ Molecular Imager^®^ and colonies were enumerated.

### Viability assays - acute treatments

For viability assays of cells acutely treated with stressors in exponential phase, wild-type and Δ*rnhC* strains were grown until OD600 0.5 at 37°C and back-diluted to OD600 0.05 in LB. For acute transcription from *P_spank(hy)_* reporters, cells were backdiluted in LB supplemented with or lacking 1mM IPTG and subsequently grown at 30°C for 2-3 generations (2 hours). For cells treated with stressors, cells were backdiluted in LB and subsequently grown at 30°C for 2-3 generations prior to the addition of the following stresses: 1.5M NaCl for 60 minutes, 1.2mM paraquat for 90 minutes, or 5μg/ml of lysozyme for 20 minutes. Cells were harvested and washed 2x with LB. After washing, samples were normalized to OD 0.3 based on the OD of the non-treated control. Cells were serially diluted and plated on LB lacking any stressor for CFU enumeration.

For viability assays of cells acutely treated with stressors in the absence of replication (stationary phase), wild-type and Δ*rnhC* strains were grown to OD600 of approximately 0.5 at 37°C and back-diluted to OD600 0.05. Cells were subsequently grown at 30°C until cells reached stationary growth. Strains were subsequently subjected to the following stress conditions: 2M NaCl for 60 minutes, 2mM Paraquat for 90 minutes, or 50 μg/ml of lysozyme for 20 minutes. For reporter strains, cells were treated with 100mM IPTG for 1 hour. Cells were harvested and washed 2x with LB. After washing, samples were normalized from the non-stress control to OD 0.3 and serial dilutions were plated for CFU enumeration.

### DNA:RNA hybrid immunoprecipitation assays (DRIPs)

DRIPs were performed as described (García-Rubio et al., 2015) with some modifications for *B. subtilis*. Precultures were diluted to OD600 of 0.05 in LB and grown at 32° C with shaking. At OD600 ~0.1, cultures were induced with 1 mM IPTG (final concentration) and grown until the culture was at OD600 = 0.3. Cells were pelleted by centrifugation and washed twice with cold PBS. Cell pellets were resuspended in spheroplast buffer (1 M sorbitol, 2 mM Tris-HCl pH 8.0, 100 mM EDTA, 0.1% β-mercapto-ethanol 0.25 mg/mL lysozyme) and incubated at 37° C for 10 minutes. Spheroplasts were collected by centrifugation at 5k RPM for 10 minutes. The supernatent was removed and pellets were washed with 1 mL of cold PBS without resuspension. Spheroplasts were then resuspended in solution I (0.8 mM GuHCl, 30 mM Tris-HCl pH 8.0, 30 mM EDTA, 5% Tween 20, 0.5% Triton X-100). Lysates were treated with RNase A (Thermo) for 30 min at 37° C and then treated with Proteinase K for 2 hours at 50° C. Lysates were then purified with chloroform-isoamyl alcohol (24:1). The aqueous phase was then moved to a new tube and 800 μL of isopropyl alcohol was added to precipitate the DNA. DNA was spooled on a Pasteur pipet and washed twice with 70% ethanol. The spooled DNA was allowed to dry at 37° C. After drying, DNA was resuspended in TE pH 8.0 and treated with HindIII overnight at 37° C. Digested chromosomal DNA was then purified over a sephadex G-50 column and brought to final volume of 125 μL. 20 μL was then removed kept as INPUT. TE was then added to the remaining sample to a final volume of 450 μL and 51 μL of 10x Binding buffer (100 mM NaPO4 pH 7.0, 1.4 M NaCl, 0.5% Triton X-100) was added. S9.6 antibody (Millipore) was added and samples were incubated overnight at 4° C with gentle rotation. After incubation with the antibody, 40 μL of 50% Protein A sepharose beads (GE) were added and IPs were incubated at 4° C for 2 hours with gentle rotation. Beads were then pelleted by centrifugation at 2000 RPM for 1 minute. The supernatant was removed and the beads were washed 3x with 1mL of 1x Binding buffer. After the washes, 120 μL of elution buffer of elution buffer II (10 mM Tris pH 8.0, 1 mM EDTA, 0.67% SDS) and 7 μL Proteinase K (Qiagen) was added. For the INPUT samples, 27 μL of TE pH 8 and 3 μL Proteinase K was added. All samples were incubated at 55° C for 45 minutes. Beads were then pelleted by centrifugation at 7000 RPMs for 1 minute and the supernatant moved to a new tube. DNA was purified using a PCR purification kit (Thermo) and analyzed using qPCR.

### Chromatin immunoprecipitation assays (ChIPs)

Precultures were diluted to OD600 of 0.05 in LB and grown at 32° C with shaking. At OD600 ~0.1, cultures were induced with 1 mM IPTG (final concentration) and grown until the culture was at OD600 = 0.3 and processed as described (Merrikh et al. 2011). Briefly, cultures were crosslinked with 1% formeldahyde for 20 minutes and subsequently quenched with 0.5 M glycine. Cell pellets were collected by centrifugation and washed once with cold phosphate buffered saline (PBS). Cell pellets were resuspended with 1.5 mL of Solution A (10 mM Tris– HCl pH 8.0, 20% w/v sucrose, 50 mM NaCl, 10 mM EDTA, 10 mg/ml lysozyme, 1mM AEBSF) and incubated at 37° C for 30 min. After incubation, 1.5 mL of 2x IP buffer (100 mM Tris pH 7.0, 10 mM EDTA, 20% triton x-100, 300 mM NaCl and 1mM AEBSF) was added and lysates were incubated on ice for 30 minutes. Lysates were then sonicated 4 times at 30% amplitude for 10 seconds of sonication and 10 seconds of rest. Lysates were pelleted by centrifugation at 8000 RPMs for 15 minutes at 4° C. Each IP was done with 1 mL of cell lysate and 40 μL was taken out prior to addition of the antibody as an input control. IPs were performed using rabbit polyclonal antibodies against DnaC (Smits et al., 2010). IPs were rotated overnight at 4° C. After incubation with the antibody, 30 μL of 50% Protein A sepharose beads (GE) were added and IPs were incubated at RT for one hour with gentle rotation. Beads were then pelleted by centrifugation at 2000 RPM for 1 minute. The supernatant was removed and the beads were washed 6x with 1 mL of 1x IP buffer. An addition wash was done with 1 mL of TE pH 8.0. After the washes, 100 μL of elution buffer I (50 mM Tris pH 8.0, 10 mM EDTA, 1% SDS) was added and beads were incubated at 65° C for 10 minutes. Beads were pelleted by centrifugation at 5000 RPMs for 1 minute. The supernatant was removed, saved and 150 μL of elution buffer II (10 mM Tris pH 8.0, 1 mM EDTA, 0.67% SDS) was added. Beads were then pelleted by centrifugation at 7000 RPMs for 1 minute and the supernatant was combined with the first elution. The combined eluates were then de-crosslinked by incubation at 65° C for overnight. The eluates were then treated with proteinase K (0.4 mg/mL) at 37° C for 2 hours. DNA was then extracted with a GeneJet PCR purification Kit (Thermo) according to the manufacturer’s instructions.

### Quantitative PCRs (qPCRs)

Quantitative PCRs (qPCRs) were performed using iTaq Universal SYBR Green master mix (Bio-Rad) and the CFX96 Touch Real-Time PCR system (Bio-Rad). For ChIPs and DRIPs, data were normalized to gene copy number by the ratios of total input to IP samples. Relative enrichment was determined by the ratio of gene copy number for *lacZ* (primers HM910/911) to *yhaX* (primers HM192/193). Relative expression was calculated as the ratio of copy number of the gene of interest (Reporter genes: *lacZ* primers HM188/189, *luxA* primers HM2799/2800, *luxE* primers HM2803/2804; Lysozyme response genes: *sigV* primers HM1766/1767, *dltA* primers HM1778/1779, *ddl* primers HM3385/3386, *murF* primers HM3387/3388; Oxidative stress response genes: *katA* primers HM3354/3355, *ahpC* primers HM3356/3357, *ykuP* primers HM3420/3421, *dhbB* primers HM3422/3423; Salt stress response genes: *proH*, primers HM3152/3153, *ysnF* primers HM3482/3483, *katE* primers HM2348/2349, *yjgC* primers HM3418/3419) to the housekeeping gene *dnaK* (primers HM770/771).

### RNA extraction and cDNA preparation

For assays with transcriptional reporters, cells were grown in LB to mid-exponential phase and backdiluted to OD600 0.05 into LB either supplemented with or lacking 1 mM IPTG. For exponential phase experiments, cells were grown for 1 hour at 37°C (3 generations) prior to harvesting. For stationary phase (replication off) experiments, cells were grown to stationary phase for 3.5 hours at 37°C. Stationary cultures were treated with IPTG in a concentration equal to 50 mM IPTG per 1 OD600 per mL of culture for one hour prior to harvesting. For assays with stress agents performed in exponential phase, mid-exponential cultures were back diluted to OD600 0.05 in LB and grown for 1 hour at 37°C (3 generations). Cultures were split and either treated or not with 0.5 μg/mL lysozyme, 0.83 M NaCl, or 0.4 mM paraquat for 10 minutes prior to harvesting. For stationary phase (replication off) experiments, cells were grown to stationary phase for 3.5 hours at 37°C. Stationary cultures were either treated or not with 10 μg/mL lysozyme for 10 minutes, 1 M NaCl for 30 minutes, or 2 mM paraquat for 120 minutes. 2-4mL of culture was harvested by addition to an equal volume of ice-cold methanol followed by centrifugation at 4,000xg for 5 minutes. Cells were lysed with 20 μg/mL lysozyme for 10 minutes for cultures grown to exponential phase, and 20 minutes for cultures grown to stationary phase. RNA was isolated with the GeneJET RNA Purification Kit (ThermoFisher Scientific). 1 μg of RNA was treated with RNase-free DNase I (ThermoFisher Scientific) for 40 minutes at 37°C. DNase I was denatured by addition of 1μl of EDTA and incubation at 65°C for 10 minutes. Reverse transcription was performed with iScript Supermix (BioRad) as per manufacturer’s instructions.

### Marker frequency

Precultures were diluted to OD600 of 0.05 in LB and grown at 32° C with shaking. At OD600 ~0.1, cultures were induced with 1 mM IPTG (final concentration) and grown until the culture was at OD600 = 0.3. Cells were pelleted by centrifugation (12k RPMs) and cell pellets were immediately frozen at -80° C. Genomic DNA was purified using a commercially available genomic DNA extraction kit (Thermo) and sent to Sinlgerla Genomics (La Jolla, CA) for DNA sequencing. Approximately 30M x 70 bp paired-end Illumina Next-Seq reads per sample were mapped to the genome of *B. subtilis* strains HM1300 (head-on lacZ) and HM1416 (co-directional lacZ) in the strain background JH642 (GenBank: CP007800.1) using Bowtie 2 (Langmead and Salzberg, 2012). Both PCR and optical duplicates were removed using Picard v1.3. Sam file was processed by SAMtools, view, sort, and mpileup functions to produce wiggle plots which were then normalized to total read count (Li et al., 2009).

### 2-dimensional gel electrophoresis

2D gels were run as described (Merrikh et al., 2015). *B. subtilis* cultures were grown to OD 0.3, then treated with 0.2% NaAzide to arrest growth. 20 mg of cells were then suspended in low-melt agarose plugs (0.5%). Lysis was performed in 2 mg/ml lysozyme for 16 hours at 37°C. Protein was removed via incubation with 5 mg/ml proteinase K, 5% sarkosyl, 0.5 M EDTA for 4 hours at 37°C. Proteinase K was then removed by 8 successive 4 hour washes in TE at 4°C. DNA was digested overnight in plugs equilibrated in 1x CutSmart buffer plus 1.0 μl of EcoRV (NEB). DNA was subjected to 2-dimensional electrophoresis and Southern blotting as previously (Friedman and Brewer, 1995). Probes for Southern blots were generated via random priming of gel-extracted PCR products corresponding to the *lacZ* region. Probes were radioactively labelled using α-32P-dATP. Hybridization images were generated with phosphor screens and imaged on a Typhoon 6000 imager (GE Healthcare).

## QUANTIFICATION AND STATISTICAL ANALYSIS

The definition of all data points and variance measurements are included in the figure legends. All measurements of standard deviation and/or standard error of means was performed in Prism 7.0 (Graphpad).

## DATA AND SOFTWARE AVAILABILITY

Deep sequencing data will be uploaded to publically accessible NCBI databases prior to publication.

## AUTHOR CONTRIBUTIONS

K.S.L., A.N.H, C.N.M, and M.R. performed experiments. K.S.L., A.N.H., C.N.M., M.R., and H.M. analyzed data. K.S.L., C.N.M., and H.M. designed experiments. K.S.L. and H.M. wrote the paper.

## ACKNOWLEDGEMENTS

We would like to thank the members of the Merrikh lab for helpful discussions and Richard Losick for his generous gift of the P*spank(hy)-luxABCDE* plasmid, pAH60. This work was supported by the National Institute of Health New Innovator Award (DP2GM110773) to H.M.

